# Asynchronous rates of lineage, phenotype, and niche diversification in a continental-scale adaptive radiation

**DOI:** 10.1101/2021.06.14.448393

**Authors:** Benjamin W. Stone, Andrea D. Wolfe

## Abstract

Rapidly diversifying clades are central to the study of diversification dynamics. This central importance is perhaps most apparent when rapid evolution occurs across several axes of diversification (*e*.*g*., lineage, phenotype, and niche); such clades facilitate investigations into the interplay between adaptive and non-adaptive diversification mechanisms. Yet, empirical evidence from rapidly evolving clades remains unclear about the relationships, if any, across diversification axes. This is especially apparent regarding the timing of diversification rate shifts. We address this knowledge gap through comparisons of the rate and timing of lineage, phenotypic, and niche diversification in *Penstemon*, a rapidly-evolving angiosperm genus. We find that diversification rate shifts in *Penstemon* are asynchronous; while we identify a burst and subsequent slowdown in lineage diversification rate ∼2.0-2.5 MYA, shifts in phenotypic and niche diversification rates either lagged behind temporally or did not occur at all. We posit that this asynchronicity in diversification rate shifts is the result of initial niche-neutral diversification followed by adaptive, density-dependent processes. Our findings contribute to a growing body of evidence that asynchronous shifts in diversification rates may be common and question the applicability of expectations for diversification dynamics across disparate empirical systems.

## Introduction

Key to the study of diversification dynamics – how and why speciation and extinction occur, and vary, across time and space – is understanding the interplay between species’ phenotypes and the environment in which they live. Of particular interest to biologists in this respect are clades which exhibit great degrees of variation in the rate of net lineage diversification (speciation minus extinction) through time, as identifying potential biotic and abiotic factors responsible for shifts in the rate of lineage diversification is integral to our understanding of the evolutionary processes generating biological diversity (Morlon, 2014). While there are numerous processes that can affect diversification dynamics, in many conceptual models, the primary sources driving diversification rate variation have an ecological basis (Aguilée et al. 2018; Aristide & Morlon, 2019).

Perhaps the most well-known of these processes is adaptive radiation: the proliferation of ecological roles and adaptations in different species within a lineage (Givnish, 1997). Adaptive radiation does not require the rapid accumulation of species *per se*, and in fact, rapid shifts in rates of diversification (‘explosive diversification’) are often incorrectly synonymized with adaptive radiation (Givnish, 2015). However, it is often the case that the two processes are linked, such that the diversification into many ecologically specialized forms via adaptive radiation also confers a sharp and temporally localized increase in the rate of lineage diversification (Gavrilets & Losos, 2009; Rundell & Price, 2009; Moen & Morlon, 2014; Martin & Richards, 2019). Integral to adaptive radiation theory is the concept of ecological opportunity, or the abundance of accessible resources underused by competing taxa (Schluter, 2000). Championed as a prerequisite for adaptive radiation by Simpson (1953), the importance of ecological opportunity stems from the idea that an abundance of accessible resources, made available through, for example, changes in environmental conditions, or the evolution of novel phenotypes, can facilitate ecological specialization and ultimately the formation of new species (Simpson, 1953; Wellborn & Langerhans, 2015; Stroud & Losos, 2016). And, like adaptive radiation, while ample ecological opportunity does not necessarily lead to an increase in clade-wide diversification rates, bursts in lineage diversification rates attributed to speciation driven by ecological opportunity are frequently observed in empirical studies (*e*.*g*., Burbrink & Pyron, 2010; Arakaki et al. 2011; García-Navas et al. 2018).

Typically, a shift in the rate of lineage diversification occurs soon after an ecological opportunity arises, producing an ‘early burst’ pattern indicative of adaptation to available niche space (Gavrilets & Losos, 2009; Yoder et al. 2010; Gillespie et al. 2020). This early burst pattern of lineage diversification can be accompanied by a burst in the rate of phenotypic diversification as well (Burbrink & Pyron, 2010; García-Navas et al. 2018). Reduced rates of phenotypic and lineage diversification are sometimes observed subsequent to early bursts (Rundell & Price, 2009; Moen & Morlon, 2014; Martin & Richards, 2019), and in cases where decreases in diversification rates are apparent, they tend to be attributed to density-dependent factors from increased competition between species as available niche space fills (Gavrilets & Vose, 2005; Ingram et al. 2012; Aguilée et al. 2018; Aristide & Morlon, 2019). However, the prevalence of diversification rate slowdowns in empirical systems, at least for phenotypic evolution, has been increasingly brought into question (Harmon et al. 2010; Slater & Friscia, 2019; Gillespie et al. 2020). Furthermore, there has been little consensus regarding the relative timing of shifts in lineage, phenotypic, and niche diversification rates. While some studies have documented close associations between the timing of lineage diversification rates and rates of phenotypic (*e*.*g*., Rabosky et al. 2013; García-Navas et al. 2018) and niche (*e*.*g*., Kozak & Wiens, 2010; Title & Burns, 2015) evolution, many others report a lack of association, or a decoupling of diversification rates (*e*.*g*., Adams et al. 2009; Folk et al. 2018; Testo & Sundue, 2018; Crouch & Ricklefs, 2019; Boucher et al. 2020). This, coupled with mixed support for the ecological opportunity hypothesis at all in continental- or global-scale radiations (*e*.*g*., Liedtke et al. 2016; Maestri et al. 2017; Folk et al. 2018; García-Navas et al. 2018), makes it unclear whether there is any general relationship between lineage, phenotypic, and niche diversification dynamics in rapidly radiating clades, especially with respect to the relative timing of rate shifts. Such uncertainty about the relationship between rates of lineage, phenotypic, and niche diversification means that expectations for a clade having potentially experienced adaptive radiation, particularly a continental radiation, are unclear. Studies of such clades, therefore, would contribute greatly to our understanding of diversification dynamics and the processes which generate biological diversity.

Here, we address this gap in our understanding of diversification dynamics through comparisons of the rate and relative timing of lineage, phenotypic, and niche diversification in a continental radiation of plants, *Penstemon* Schmidel (Plantaginaceae). Commonly known as the beardtongues, the genus *Penstemon* has nearly 300 described species, and represents the most speciose genus of angiosperms endemic to North America (Wolfe et al. 2006; Freeman, 2019). *Penstemon* exhibits an exceptional array of phenotypic diversity in floral and vegetative characters, with the degree floral diversity in particular suggesting a substantial history of selective pressure from pollinators (Straw, 1966). Species of *Penstemon* occupy a wide variety of ecological niches, with several examples of edaphic specialization (*e*.*g*., sand dunes, oil shales, and calcareous soils), although most species prefer semi-disturbed, xeric habitats (Wolfe et al. 2006; Wolfe et al. 2021). This tendency toward habitat specialization and the limited geographic distributions that often accompany it has resulted in conservation concerns for many *Penstemon* species (*e*.*g*., Wolfe et al. 2014, 2016; Rodriguez-Peña et al. 2018; Stone et al. 2019, 2020; Zacarías-Correa et al. 2020). Phylogenetic studies of *Penstemon* have revealed diversification patterns consistent with a rapid evolutionary radiation, including difficulty with estimating relationships between clades early in the evolutionary history of the genus, likely due to the effects of incomplete lineage sorting (Wolfe et al. 2006; Wessinger et al. 2016, 2019), and an early, rapid burst in the rate of lineage diversification coincident with glacial activity during the Pleistocene (Wolfe et al. 2021). Consequently, the degree of phenotypic diversity and niche specialization, and the presence of an early burst pattern of lineage diversification coincident with environmental changes at a global scale, has resulted in the hypothesis that *Penstemon* represents a continental example of a rapid adaptive radiation (Wolfe et al. 2006, 2021). However, to date, no analysis of the relative timing of shifts in lineage, phenotypic, and niche diversification has been conducted in the context of the *Penstemon* radiation.

We built data sets of ecological niche and phenotypic trait variables, and, using a modified version of the time-calibrated *Penstemon* phylogeny inferred in Wolfe et al. (2021), analyzed macroevolutionary patterns of lineage, phenotypic, and niche diversification in *Penstemon*. Using these data, we aim to answer the following questions: (1) Do shifts in the rates of phenotypic and niche diversification coincide with the shift in lineage diversification rate? (2) Do rates of lineage diversification depend on the states of phenotypic or environmental variables? (3) Does the timing of diversification coincide with potential increases in ecological opportunity due to changes in glacial activity and global temperature?

## Methods

### Data generation

The phylogeny used in this study is a modified version of the time-calibrated phylogeny inferred in Wolfe et al. (2021). Briefly, this phylogeny was constructed from 43 nuclear loci generated from a targeted amplicon sequencing approach described in Blischak et al. (2014), and inferred with *RAxML* v8.2.10 (Stamatakis, 2014). Divergence times were calibrated with previously inferred divergence events in the Lamiales (Vargas et al. 2014) in *BEAST* v2.6.0 (Bouckaert et al. 2019) using a relaxed log-normal clock model (Drummond et al. 2006); these estimates were then used as secondary calibration bounds to date the phylogeny using *treePL* (Smith & O’Meara, 2012). More details about the methodology used to produce the time-calibrated *Penstemon* phylogeny can be found in Wolfe et al. (2021).

Species occurrence data were collected for each species of *Penstemon* that was both in the constructed phylogeny and available on GBIF (gbif.org). GBIF coordinates were downloaded on June 26, 2020. For ten species (listed in Supplemental Table 1), we added coordinates from the primary literature and from herbarium records accessed through the SEINet portal (swbiodiversity.org/seinet/). We retained all occurrence points accurate to at least three decimals for both latitude and longitude, and removed occurrences which had identical coordinates or were clearly outside the species’ described range. Finally, we included only taxa for which there were at least three occurrences in the data set following these curation measures.

To build the data set to evaluate the evolution of ecological niche space, we first extracted 28 environmental layers capturing features of the climate, soil, landcover, and topology at 30s resolution. These layers included: 19 temperature and precipitation variables (bioclim variables from worldclim.org), elevation (from worldclim.org), slope, aspect, and six landcover classes from earthenv.org/landcover. Values for slope and aspect were calculated in R, using the elevation raster file. The six landcover classes correspond to the percentage landcover of evergreen/deciduous needleleaf trees, evergreen broadleaf trees, deciduous broadleaf trees, mixed/other trees, shrubs, and herbaceous vegetation. For each GBIF occurrence point, environmental conditions were extracted in R, and for each taxon, median values of each predictor were used in downstream analyses. Environmental data were then ordinated via phylogenetic principal components analysis (pPCA) using the ‘phyl.pca’ function on the correlation matrix in the R package *phytools* (Revell, 2012), forming a composite variable summarizing species’ niche space. All downstream analyses using both the niche data set and the phylogeny were performed with a pruned phylogeny including only taxa for which niche data were collected.

Phenotypic traits for species were collected primarily from species descriptions in the Flora of North America (Freeman, 2019). For species not included in the Flora (mainly those endemic to Mexico), we accrued data through a combination of primary species’ descriptions and measurements of specimens from herbarium collections. For these species (listed in Supplemental Table 1), we downloaded images of herbarium specimens via the SEINet portal, the Steere Herbarium at the New York Botanical Gardens, and the National Autonomous University of Mexico (UNAM) Herbarium and made virtual measurements of focal phenotypic characters with *ImageJ* (Schneider et al. 2012). We collected both categorical and continuous traits: the full list of traits can be found in Supplemental Table 2. For continuous traits, we recorded the midpoint value, after outlier measurements were discarded. For categorical traits, we coded each descriptor sequentially, starting from zero. For species with multiple or ambiguous categorical characters, a single trait value was selected at random. We used the R package *StatMatch* (D’Orazio, 2012) to compute a phenotypic distance matrix from Gower’s Distance metric (Gower, 1971), calculated with the correction proposed by Kaufman and Rousseeuw (1990). All downstream analyses using both the phenotypic data set and the phylogeny were performed with a pruned phylogeny including only taxa for which phenotypic data were collected. To ordinate the phenotypic data, we conducted generalized multidimensional scaling (PCOA) on the distance matrix using the ‘pcoa’ function in the R package *ape* (Paradis & Schliep, 2019), using the Cailliez correction (Cailliez, 1983). Because PCOA does not produce traditional loadings in the same sense that PCA does, we produced an alternative measure of variable importance for each PCOA axis. We did so by regressing the first axis of the ordinated data with each uncorrected variable and dividing the R^2^ for each variable by the sum of all R^2^ values, producing a relative measure of variable importance. We also constructed these values for the niche data set, to facilitate comparisons of niche and phenotype variable importance.

### Ancestral state reconstruction and phylogenetic generalized least-squares

We performed ancestral state reconstruction on the ordinated phenotypic and niche data sets using the *phylopars* package (Bruggeman et al. 2009) in R. We fit models of continuous trait evolution under Brownian motion (BM), Ornstein-Uhlenbeck (OU) and Early Burst (EB) models, and chose the best model using the corrected Akaike information criterion (AICc; Akaike, 1973). Then, using the reconstruction from the best model, we calculated the degree of the trait shift at each node by finding the absolute difference between the reconstructed values at a focal node and its parent node. Major shifts were identified as values in the 95^th^ percentile of trait shifts.

To investigate evolutionary associations between ecological niche space and phenotype, we performed phylogenetic generalized least squares regression (PGLS). PGLS generates slope and intercept estimates which account for interspecific autocorrelation of residuals due to species’ shared phylogenetic histories. Because our goal was to evaluate the potential impact of environmental predictor variables on the phenotype, we conducted all PGLS analyses with the first axis of the phenotype PCOA (phenotype summary statistic) as the response variable, and for predictor variables, we used the first axis of the niche pPCA (niche summary statistic) and individual uncorrected environmental variables. For each predictor variable, we performed PGLS under Pagel’s *λ* model, which is a generalization of the assumption of Brownian Motion (Pagel, 1997). In this model, *λ*can range from 0 to 1; when *λ* = 0, the analysis is equivalent to ordinary least squares regression, and when *λ* = 1, the analysis is equivalent to phylogenetic independent contrasts (residual errors are Brownian). All PGLS analyses were conducted with the *nlme* package in R (Pinheiro et al. 2021).

### Macroevolutionary rates analyses

We used *BAMM* (Rabosky, 2014) to assess rates of net lineage diversification through time. Prior settings were derived from the R package *BAMMtools* (Rabosky et al. 2014). We ran the *BAMM* analysis for 50 million MCMC steps, in four chains, sampling every 10,000 steps. We set the minimum clade size for shift inference equal to two (no terminal branch shifts), and set the global sampling fraction to 0.9. Otherwise, parameter values were left at their default values, or were inferred from *BAMMtools*. During post-processing, the first 10% of samples were discarded as burn-in. We also used the *BAMM* trait model to assess the rate of both phenotypic and niche diversification through time. The first axes of the ordinated phenotypic and niche data sets were used as input, and we used *BAMMtools* to generate prior settings as with the diversification analysis. For the phenotypic data, we ran the analysis for 2 billion steps, sampling every 100,000 steps, and for the niche data, we ran the analysis for 250 million steps, sampling every 10,000 steps. The first 50% and 10% of samples for the phenotypic and niche data, respectively, were discarded as burn-in during post-processing. All other settings followed those described for the *BAMM* diversification analysis. To examine variation in rates of diversification in the context of particular clades, we also generated rate-through-time plots for eleven clades on the basis of their importance in the context of *Penstemon* taxonomy.

We used a regression approach to assess potential relationships between macroevolutionary rates and global temperature. To do so, we first obtained global temperature estimates over the past ∼3.5 million years, which were derived from a temperature anomaly data set (de Boer et al. 2014). We then generated a *BAMM* rates-through-time matrix for each of the three *BAMM* analyses, obtaining estimates of the average rate diversification metric at 100 discrete points in time (roughly 1000-year time slices). Values for the rate diversification metric were then paired with global temperature estimates to the nearest 1,000-year interval. We then performed regression analyses with linear, exponential, and quadratic regression, and selected the best model for each diversification metric using AIC.

We used *ES-sim* (Harvey & Rabosky, 2018) to test for relationships between trait variation and variance in lineage diversification. We performed this test for both phenotypic and niche data sets, using the first axis of each ordinated data set as input. We also ran *ES*-*sim* on each individual continuous phenotypic and environmental trait, to test relationships between a particular trait and rates of lineage diversification.

## Results

### Data generation

In total, our data collection and processing efforts generated a phenotypic data set for 280 taxa and a niche data set for 277 taxa (from 14,495 GBIF occurrences). The taxa included in both data sets are identical, except for three taxa (*P. debilis, P. wendtiorum*, and *P. tracyi*) not included in the niche data set because of a lack of GBIF occurrences owed to the rarity of these species and (for *P. debilis* and *P. tracyi*) their protected status due to conservation concerns. Relative measures of variable importance for the ordinated phenotypic and niche data sets can be found in Tables 1 and 2, respectively. The phenotypic traits with a corrected R^2^ > 0.05 (analogous to loadings in a traditional PCA) were inflorescence type, leaf length, flower presentation, lower topography, throat expansion, plant form, and the presence/absence of basal leaves (Table 1). For the niche data set, variables meeting this criterion were mostly measures of temperature and precipitation seasonality, and the percentage of herbaceous vegetation (Table 2). Species with higher values for the niche summary statistic tend to be at higher elevations with more herbaceous landcover, more temperature seasonality (but lower minimum, maximum, and mean temperatures), and less precipitation seasonality (with less overall precipitation, but comparatively more precipitation during the dry season).

**Table 1.**
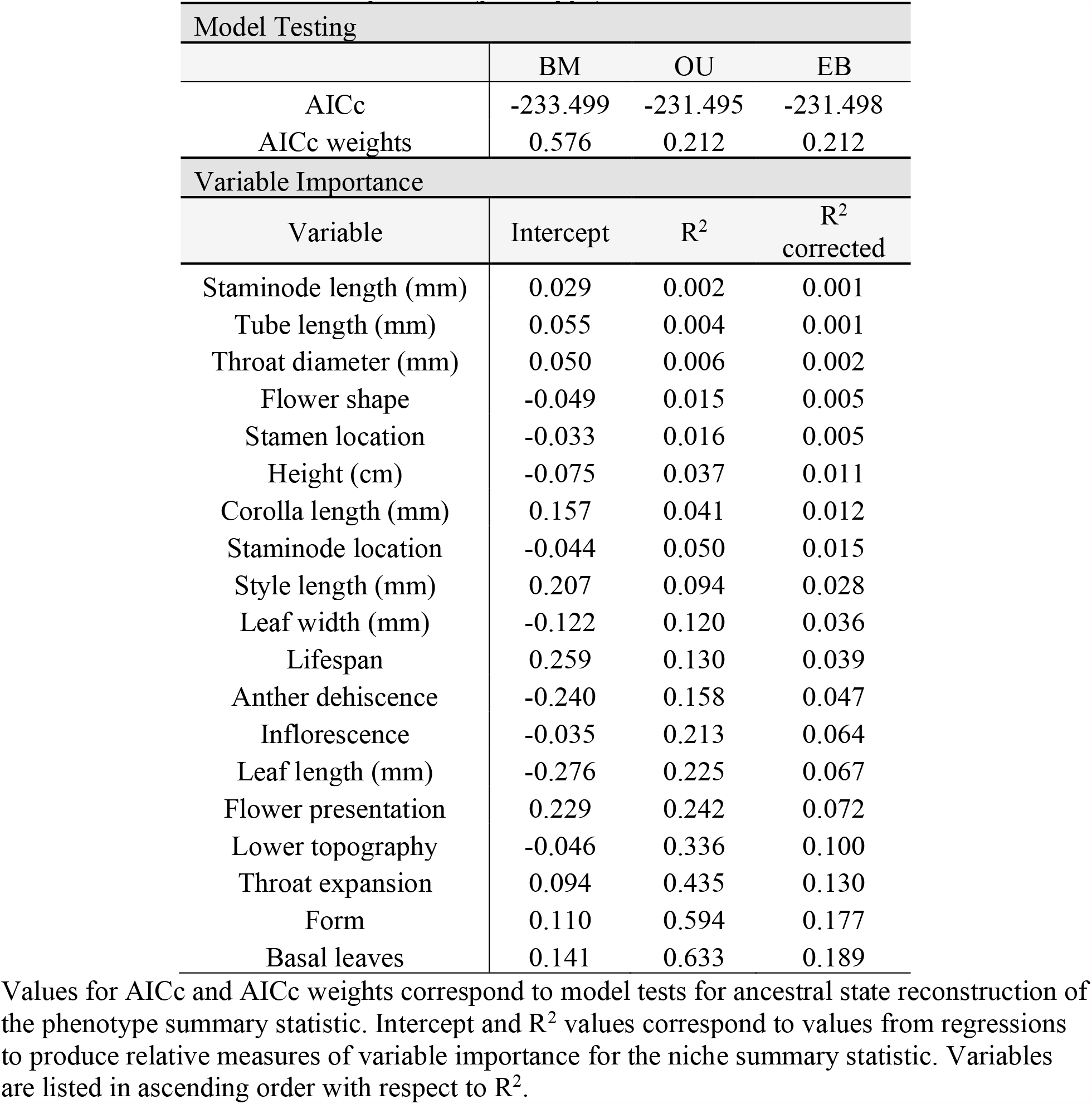
Model weights for Ancestral State Reconstruction and variable importance (phenotype)

| Model Testing |  |  |  |
| --- | --- | --- | --- |
|  | BM | OU | EB |
| AICc | -233.499 | -231.495 | -231.498 |
| AICc weights | 0.576 | 0.212 | 0.212 |
| Variable Importance |  |  |  |
| Variable | Intercept | R<sup>2</sup> | R<sup>2</sup><br>corrected |
| Staminode length (mm) | 0.029 | 0.002 | 0.001 |
| Tube length (mm) | 0.055 | 0.004 | 0.001 |
| Throat diameter (mm) | 0.050 | 0.006 | 0.002 |
| Flower shape | -0.049 | 0.015 | 0.005 |
| Stamen location | -0.033 | 0.016 | 0.005 |
| Height (cm) | -0.075 | 0.037 | 0.011 |
| Corolla length (mm) | 0.157 | 0.041 | 0.012 |
| Staminode location | -0.044 | 0.050 | 0.015 |
| Style length (mm) | 0.207 | 0.094 | 0.028 |
| Leaf width (mm) | -0.122 | 0.120 | 0.036 |
| Lifespan | 0.259 | 0.130 | 0.039 |
| Anther dehiscence | -0.240 | 0.158 | 0.047 |
| Inflorescence | -0.035 | 0.213 | 0.064 |
| Leaf length (mm) | -0.276 | 0.225 | 0.067 |
| Flower presentation | 0.229 | 0.242 | 0.072 |
| Lower topography | -0.046 | 0.336 | 0.100 |
| Throat expansion | 0.094 | 0.435 | 0.130 |
| Form | 0.110 | 0.594 | 0.177 |
| Basal leaves | 0.141 | 0.633 | 0.189 |

Values for AICc and AICc weights correspond to model tests for ancestral state reconstruction of the phenotype summary statistic. Intercept and R<sup>2</sup> values correspond to values from regressions to produce relative measures of variable importance for the niche summary statistic. Variables are listed in ascending order with respect to R<sup>2</sup>.

**Table 2.** Model weights for Ancestral State Reconstruction and variable importance (niche)

| Model Testing |  |  |  |
| --- | --- | --- | --- |
|  | BM | OU | EB |
| AICc | 1402.215 | 1404.219 | 1401.994 |
| AICc weights | 0.403 | 0.148 | 0.450 |
| Variable Importance |  |  |  |
| Variable | Intercept | R<sup>2</sup> | R<sup>2</sup> corrected |
| Slope | 1.016 | 0.001 | 0.000 |
| Aspect | 1.236 | 0.001 | 0.000 |
| % shrubs | 0.895 | 0.002 | 0.000 |
| Precip. of driest quarter | 0.299 | 0.006 | 0.001 |
| % deciduous broadleaf trees | 0.844 | 0.012 | 0.002 |
| % evergreen broadleaf trees | 0.783 | 0.014 | 0.002 |
| Precip. of driest month | -0.057 | 0.025 | 0.003 |
| Mean temp. of wettest quarter | 1.777 | 0.038 | 0.005 |
| Mean diurnal temp. range | -7.644 | 0.049 | 0.006 |
| % mixed/other trees | 1.757 | 0.050 | 0.006 |
| % needleleaf trees | 1.564 | 0.051 | 0.007 |
| Max temp. of warmest month | 8.354 | 0.062 | 0.008 |
| Precip. of warmest quarter | 2.654 | 0.108 | 0.014 |
| Mean temp. of warmest quarter | 8.376 | 0.168 | 0.022 |
| elevation | -3.411 | 0.171 | 0.022 |
| Precip. of coldest quarter | 2.562 | 0.204 | 0.026 |
| Annual precip. | 4.372 | 0.297 | 0.038 |
| Mean temp. of driest quarter | 3.815 | 0.433 | 0.056 |
| Precip. of wettest quarter | 4.465 | 0.452 | 0.058 |
| % herbaceous vegetation | -2.176 | 0.458 | 0.059 |
| Precip. of wettest month | 4.683 | 0.472 | 0.061 |
| Isothermality | 18.126 | 0.506 | 0.065 |
| Annual mean temp. | 6.886 | 0.564 | 0.073 |
| Precip. seasonality | 7.004 | 0.583 | 0.075 |
| Temp. seasonality | -13.212 | 0.652 | 0.084 |
| Annual temp. range | -22.126 | 0.696 | 0.090 |
| Mean temp. of coldest quarter | 0.926 | 0.809 | 0.104 |
| Min temp. of coldest month | -3.869 | 0.865 | 0.112 |

Values for AICc and AICc weights correspond to model tests for ancestral state reconstruction of the niche summary statistic. Intercept and R<sup>2</sup> values correspond to values from regressions to produce relative measures of variable importance for the niche summary statistic. Variables are listed in ascending order with respect to R<sup>2</sup>.

### Ancestral state reconstruction and phylogenetic generalized least-squares

The best ancestral state reconstruction model for the ordinated phenotypic data set was the BM model, which had an AICc score slightly more than 2 units less than the next closest models and comprised over half of the model weight (Table 1). Our method for identifying shifts in ancestral phenotype indicated that two of the three largest shifts in phenotype occur at the base of clades containing subgenus *Dasanthera* and subgenus *Penstemon* section *Caespitosi* (Figure 1). These two clades also occupy a similar phenotypic space with respect to the ordinated phenotypic data set. Values of the phenotype summary statistic are similar between these species, representing some of the lowest values in the genus (Figure 1). This reflects the fact that species in these clades tend to exhibit a shrub or subshrub growth form with no basal leaves, and with corolla throats expanded on the upper side. These characters were determined to be among the most important to the PCOA (Table 1), making it unsurprising that species with similarities in these traits have similar values for the phenotype summary statistic.

**Figure 1.**
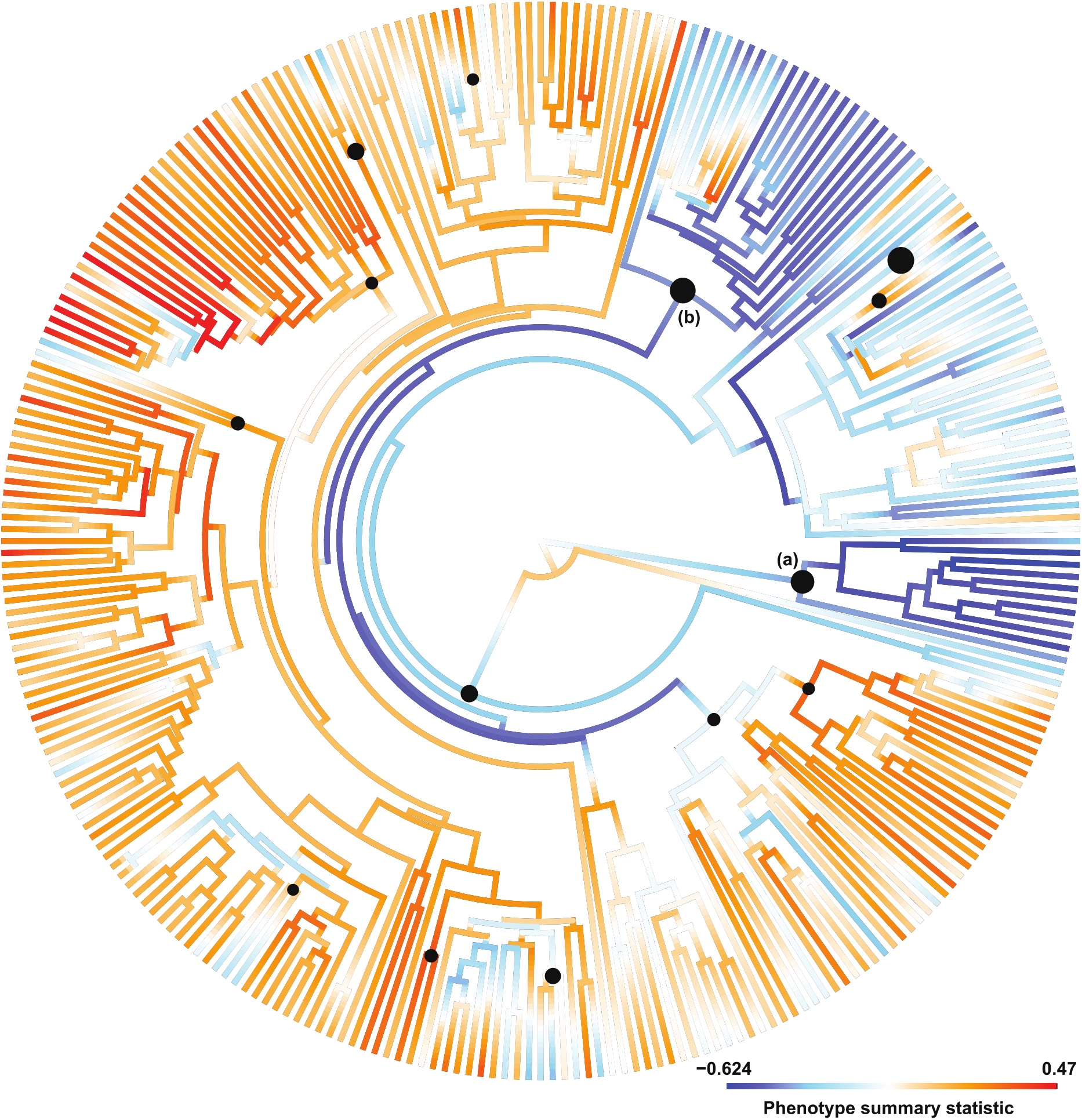
Ancestral state reconstruction of the phenotype summary statistic. Labeled clades contain (a) subgenus *Dasanthera* and (b) section *Caespitosi*. Trait values represent scores of the first PC axis of ordinated phenotype data; cooler values indicate (generally) short plants with a shrub or subshrub growth form, no basal leaves, and with corolla throats expanded on the upper side. Black circles at nodes indicate the largest shifts (top 5%) in ancestral state estimates from the preceding node. Circles are scaled to the size of the shift in trait value.

For the niche data set, the best model for ancestral state reconstruction was the EB model (Table 2). However, this model was nearly indistinguishable from the BM model, with AICc scores for both models very similar. Both the BM and EB models had AICc scores close to 2 units less than the OU model, and together, they comprised > 85% of the total model weight. Some of the largest shifts in ancestral niche space were located at the base of clades containing subgenus *Penstemon* sections *Fasciculus* and *Caespitosi* (Figure 2). This corresponds with a shift in geographic distribution for section *Fasciculus*; the species in this clade are among the most southernly-distributed *Penstemon* species, found almost exclusively in Mexico and Guatemala. The ancestral state reconstruction reflects this shift to habitats at lower elevations with less temperature variability (but higher overall temperatures) and more precipitation variability (with more overall precipitation, except during the dry season). There are a few other clade-wide shifts (*e*.*g*., at the base of section *Caespitosi*), but these may be artifacts produced by the ancestral state reconstruction method, such that large shifts in the trait estimate at a given node does not necessarily relate to the estimate of the trait along the terminal branch. In the case of section *Caespitosi*, a shift from a positive to a negative value occurs at the node in question, but the majority of taxa in this clade exhibit a positive value for the niche summary statistic (Figure 2).

**Figure 2.**
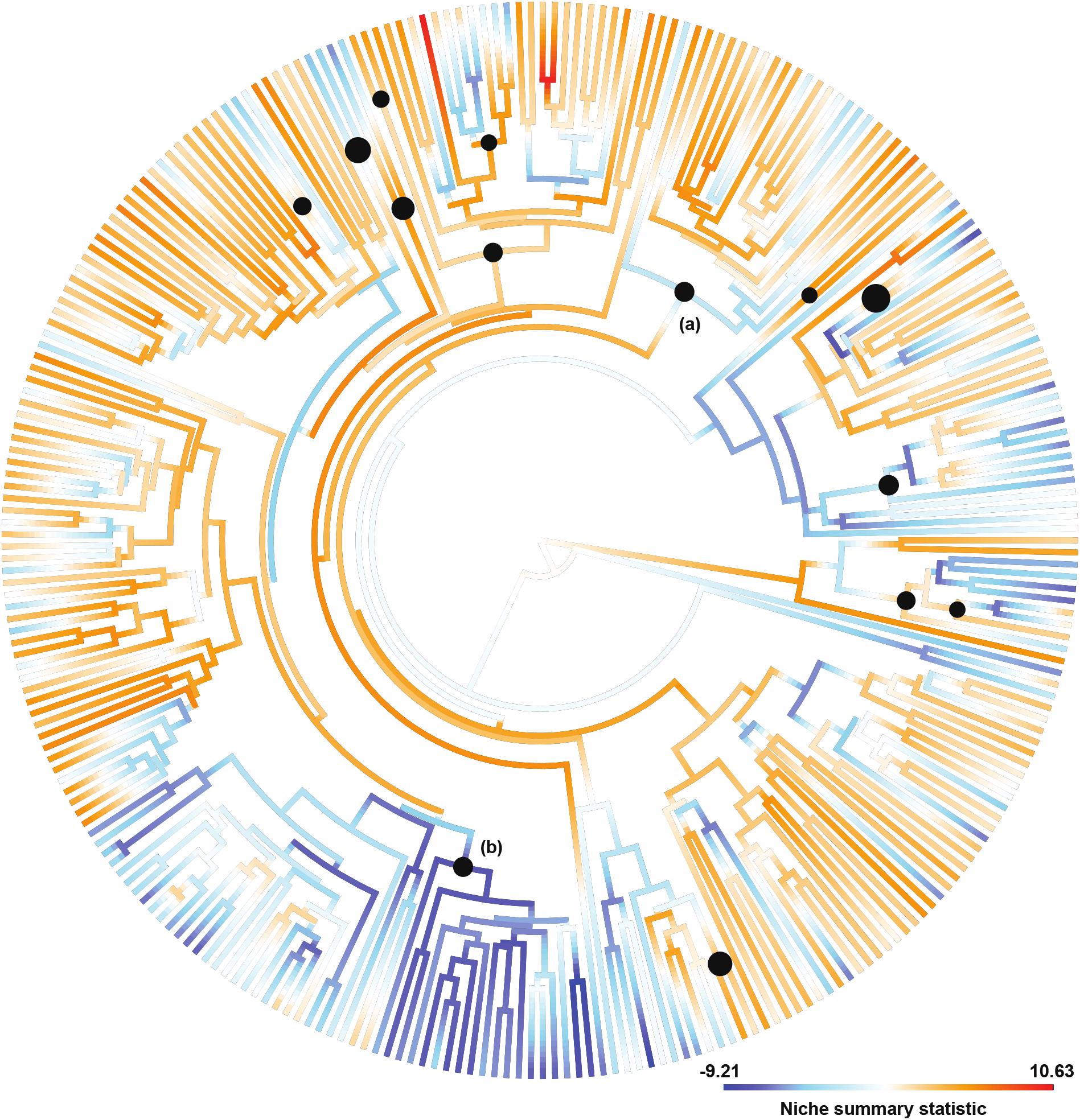
Ancestral state reconstruction of the niche summary statistic. Labeled clades contain sections (a) *Caespitosi* and (b) *Fasciculus*. Trait values represent scores of the first PC axis of ordinated niche data; warmer values indicate (generally) higher elevations with more herbaceous landcover, more temperature seasonality (but lower minimum, maximum, and mean temperatures), and less precipitation seasonality (with less overall precipitation, but comparatively more precipitation during the dry season). Black circles at nodes indicate the largest shifts (top 5%) in ancestral state estimates from the preceding node. Circles are scaled to the size of the shift in trait value.

For each PGLS analysis, estimates of *λ* were high, with all point estimates of *λ* > 0.95 and confidence intervals capped at the upper parameter bound (Table 3). This suggests a strong phylogenetic signal to the data, such that the shared phylogenetic history between species strongly affects the error structure of the residuals, and that this error structure is similar to expectations under Brownian motion. Only two environmental variables had a significant effect on the phenotype summary statistic, suggesting that there is no significant relationship between the residuals of the regression between the phenotype summary statistic and the majority of the environmental variables tested, after accounting for expected phylogenetic covariance (Table 3). The two variables with significant effects were the percentage of evergreen broadleaf trees (*λ* = 0.987, slope = -1.05 x 10^−2^, *p* = 0.001) and the amount of precipitation during the warmest quarter (*λ* = 0.983, slope = -3.05 x 10^−4^, *p* = 0.038).

**Table 3.** PGLS results

| Variable | $\lambda$ | 95% confidence interval | Slope | *p*-value |
| --- | --- | --- | --- | --- |
| Niche summary statistic | 0.959 | (0.868, 1) | 1.45E-03 | 0.634 |
| Elevation | 0.960 | (0.871, 1) | 9.96E-06 | 0.547 |
| Aspect | 0.962 | (0.873, 1) | -1.33E-04 | 0.377 |
| Slope | 0.962 | (0.873, 1) | 1.81E-03 | 0.720 |
| % needleleaf trees | 0.962 | (0.873, 1) | 5.46E-04 | 0.289 |
| **% evergreen broadleaf trees** | 0.987 | (0.901, 1) | -1.05E-02 | **0.001** |
| % deciduous broadleaf trees | 0.965 | (0.876, 1) | -5.11E-03 | 0.165 |
| % mixed/other trees | 0.965 | (0.877, 1) | 7.97E-04 | 0.350 |
| % shrubs | 0.961 | (0.872, 1) | -2.27E-04 | 0.586 |
| % herbaceous vegetation | 0.960 | (0.870, 1) | 2.69E-04 | 0.691 |
| Annual mean temp. | 0.958 | (0.867, 1) | -1.12E-03 | 0.668 |
| Mean diurnal temp. range | 0.963 | (0.873, 1) | -8.86E-03 | 0.180 |
| Isothermality | 0.954 | (0.861, 1) | -1.64E-03 | 0.420 |
| Temp. seasonality | 0.961 | (0.870, 1) | -3.04E-07 | 0.997 |
| Max temp. of warmest month | 0.959 | (0.869, 1) | -2.23E-03 | 0.444 |
| Min temp. of coldest month | 0.961 | (0.870, 1) | -7.30E-05 | 0.972 |
| Annual temp. range | 0.965 | (0.875, 1) | -1.17E-03 | 0.629 |
| Mean temp. of wettest quarter | 0.962 | (0.873, 1) | 5.13E-04 | 0.747 |
| Mean temp. of driest quarter | 0.962 | (0.872, 1) | 2.26E-04 | 0.867 |
| Mean temp. of warmest quarter | 0.959 | (0.868, 1) | -1.70E-03 | 0.556 |
| Mean temp. of coldest quarter | 0.958 | (0.866, 1) | -8.21E-04 | 0.702 |
| Annual precip. | 0.963 | (0.873, 1) | -1.41E-05 | 0.692 |
| Precip. of wettest month | 0.963 | (0.873, 1) | -1.18E-04 | 0.550 |
| Precip. of driest month | 0.961 | (0.870, 1) | 7.95E-05 | 0.936 |
| Precip. seasonality | 0.962 | (0.872, 1) | -6.58E-04 | 0.212 |
| Precip. of wettest quarter | 0.963 | (0.873, 1) | -3.91E-05 | 0.577 |
| Precip. of driest quarter | 0.962 | (0.872, 1) | -4.59E-05 | 0.879 |
| **Precip. of warmest quarter** | 0.983 | (0.896, 1) | -3.05E-04 | **0.038** |
| Precip. of coldest quarter | 0.961 | (0.871, 1) | -4.87E-06 | 0.950 |

All PGLS analyses were conducted with the phenotype summary statistic as the dependent variable. Environmental variables listed in the first column constitute independent variables. $\lambda$ can range from 0 to 1; at $\lambda = 0$ , the model simplifies into ordinary least squares regression, and at $\lambda = 1$ , the model simplifies into phylogenetic independent contrasts. Significant *p*-values ( $p < 0.05$ ) are indicated in bold and correspond to the significance of the slope after accounting for expected covariance due to the phylogeny.

### Macroevolutionary rates analyses

For all three evolutionary rates analyses in *BAMM*, neither the estimates of effective sample size (ESS) or visualization of trace plots suggested failed MCMC chain convergence. For the lineage diversification analysis, the best model was one with a single rate shift (Supplemental Table 3). Of the credible set of shift configurations, the single best shift configuration (proportion of posterior samples, *f*, = 0.88) suggests a single burst in the rate of lineage diversification about 2.0-2.5 MYA (Figure 3a). This shift includes all major clades of *Penstemon* except subgenus *Dasanthera, P. personatus, P. scapoides*, and *P. caesius*. Subsequent to this burst, the rate of diversification slowed continuously to the present day, where estimates of speciation rates are the lowest (Figure 3b). Our results very closely resemble those found in Wolfe et al. (2021) and are consistent with the hypothesis that *Penstemon* represents a recent and rapid evolutionary radiation (Wolfe et al. 2006).

**Figure 3.**
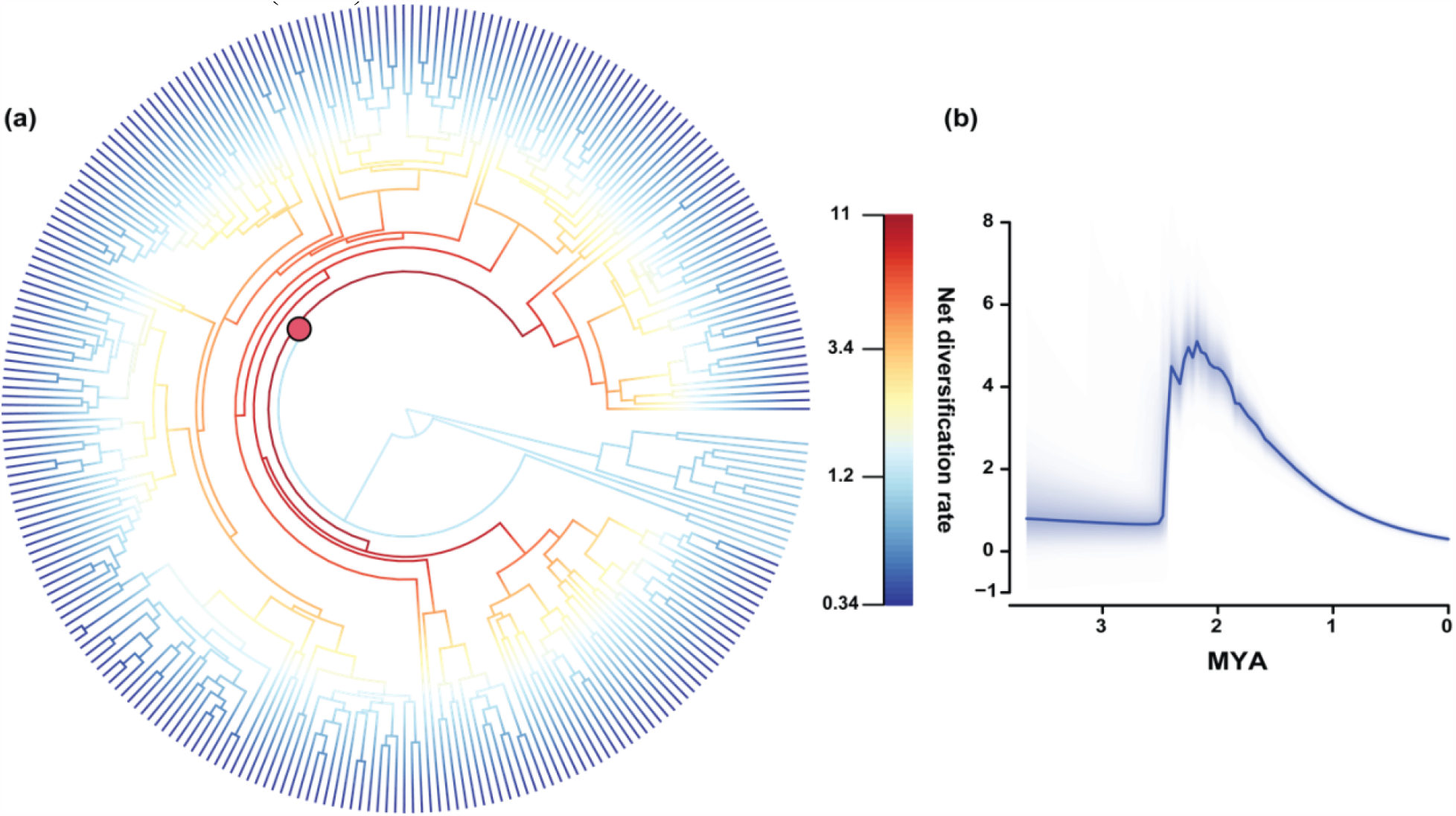
Best shift configuration (a) and rates-through-time (RTT) plot (b) of *Penstemon* lineage diversification rates in *BAMM*. (a) *BAMM* suggested a single shift in the rate of lineage diversification early in the evolutionary history of genus (red dot), followed by a subsequent slowdown of rates of lineage diversification. (b) Lineage diversification rates increased sharply between 2.0-2.5 MYA, declining steadily thereafter. Both findings are consistent with results from Wolfe et al. (2021).

Analyses of rate shifts in phenotypic diversification were inconclusive, with Bayes Factors continuing to increase with larger numbers of shifts, and posterior probabilities spread across shift configurations (Supplemental Table 4). Plots of clade-specific diversification rates show that sections *Fasciculus* and *Spectabiles* exhibit a burst in phenotypic diversification, while section *Caespitosi* shows a slowdown of phenotypic diversification (Supplemental Figure 1). Analyses of niche diversification were more informative. Although Bayes Factors continued to increase as the number of shifts increased, posterior probabilities place most weight on 0-3 shifts with a posterior probability of 0.42 for zero shifts and 0.28 for a single shift affecting two species (Supplemental Table 5). Plots of clade-specific diversification rates did not reveal differences between focal taxonomic groups, suggesting that our data do not point clearly to any particular shifts in niche diversification rate, although the presence of such shifts cannot be ruled out (Supplemental Figure 1).

Regressions between global temperature and lineage diversification rate indicated the quadratic model as the best fit, with > 99% of the AIC model weight, and a correlation coefficient (Kendall’s *τ*) of 0.33 (Supplemental Table 6). This model suggests that lineage diversification rates were highest at intermediate global temperatures (Figure 4). The regressions between global temperature and phenotypic diversification produced somewhat equivocal results; the exponential and quadratic regressions gave very similar AIC scores, with roughly equal model weights for both, and correlation coefficients (*τ*) were also very similar (Supplemental Table 6). However, plots of both models support the same positive correlation between global temperature and phenotypic diversification rates. Regressions between global temperature and niche diversification were similar to those for phenotypic diversification; the exponential model is a slightly better fit than the quadratic model, and the correlation coefficients (*τ*) are very similar (Supplemental Table 6), but plots of both models suggest a positive correlation between global temperature and niche diversification rates. We therefore present only the best models for the phenotypic and niche regression analyses (Figure 4), despite only miniscule differences between the best and second-best models.

**Figure 4.**
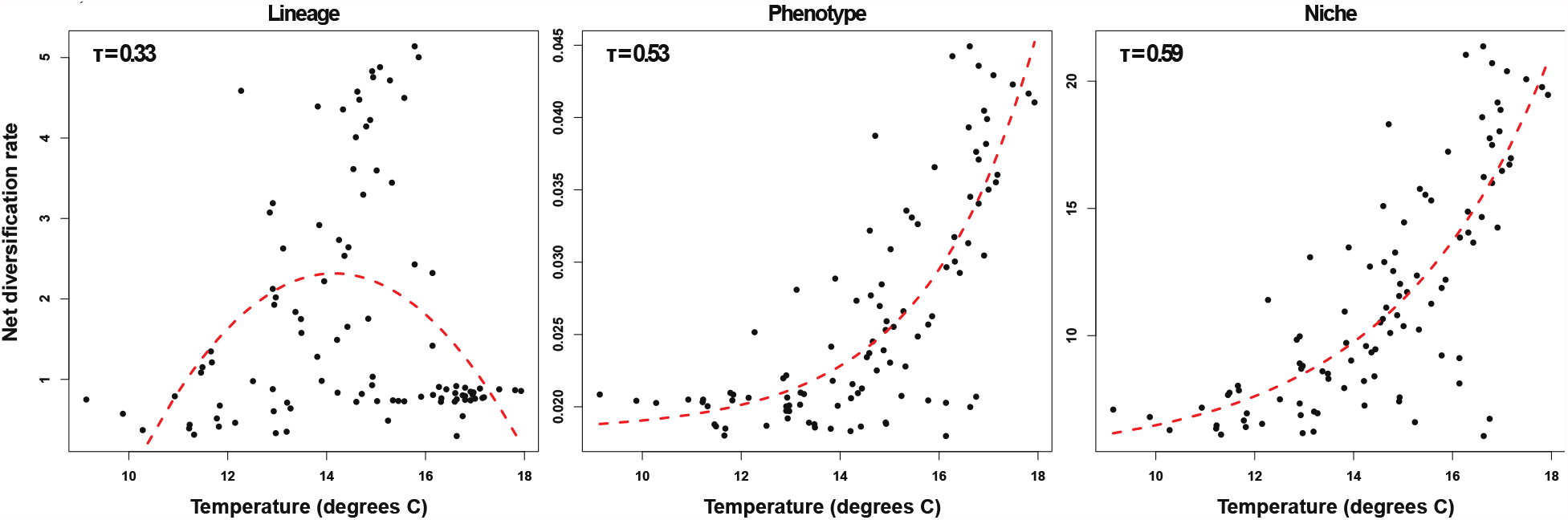
Best regressions of global temperature with diversification rates. Global temperatures through time were inferred from climate model simulations of benthic delta-O-18 (de Boer et al. 2014).

Results of all significant *ES-sim* analyses can be found in Table 4. These analyses did not reveal statistically significant relationships between variance in either the phenotype or niche summary statistic and variance in rates of lineage diversification. However, tests on unordinated phenotypic and environmental variables produced significant results for plant height, annual mean temperature (bioclim variable 1), and mean temperature of the coldest quarter (bioclim variable 11). These results suggest a correlative relationship between mean temperature, plant height, and the rate of lineage diversification, such that taller plants, and warmer annual and winter temperatures, lead to higher rates of lineage diversification.

**Table 4.** Significant *ES-sim* results

| Variable | $\rho$ | R<sup>2</sup> | *p*-value |
| --- | --- | --- | --- |
| Annual mean temp. | 0.307 | 0.094 | 0.039 |
| Mean temp. of coldest quarter | 0.296 | 0.088 | 0.047 |
| Height (cm) | 0.351 | 0.123 | 0.015 |

$\rho$ is the population Pearson correlation coefficient.

## Discussion

We analyzed macroevolutionary patterns of lineage, phenotypic, and niche diversification in *Penstemon*, a large angiosperm genus that has recently undergone a continental-scale rapid evolutionary radiation. Our results indicate that although *Penstemon* experienced an early burst of lineage diversification rates, the peak of which occurred approximately 2.0-2.5 MYA (Figure 3), there is no clear evidence that this was accompanied by an increase in the rate of phenotypic or niche evolution. Conversely, it appears that changes in phenotypic diversification rates likely operate at a smaller taxonomic scale, changing in smaller clades, rather than genus-wide, and occurring well after the initial burst in lineage diversification rates (Supplemental Figure 1). We did, however, find evidence for higher rates of lineage diversification in warmer environments and in larger plants (Table 4), and it appears that, generally, rates of lineage, phenotypic, and niche diversification are correlated with global temperature (Figure 4). Our results contribute to a growing body of evidence suggesting that asynchronicity in diversification rate shifts may be common, and bring into question the general applicability of expectations for diversification dynamics, which are derived mainly from studies of island systems, to studies of continental radiations.

### Asynchronous shifts in rates of lineage, phenotypic, and niche diversification

We observed a distinct tree-wide shift in the rate of net lineage diversification approximately 2.0-2.5 MYA (Figure 3) which is not evident in phenotypic or niche diversification. Rather, for niche evolution, most evidence points to no shifts in diversification rate at all (Supplemental Figure 1, Supplemental Table 5). For phenotype evolution, although the exact number of rate shifts is inconclusive (Supplemental Table 4), no single coincident shift is apparent, suggesting that changes in phenotypic diversification rate occur at a smaller taxonomic scale (intra-clade vs. entire genus) and lag temporally behind the shift in lineage diversification rate (Supplemental Figure 1). An important caveat about our results is that our summarizations of diversity in environmental and phenotypic space in *Penstemon* are not comprehensive. There are potentially important unmeasured phenotypic and ecological variables that, if added to our analyses, may indeed produce synchronous diversification rate shifts. Examples of this include floral volatiles integral to pollinator attraction (Parachnowitsch et al. 2012), interactions with arbuscular mycorrhizae (Titus & del Moral, 1998), and a diverse array of secondary metabolites for defense against herbivory (*e*.*g*., Kelly & Bowers, 2016, 2018). We highlight this caveat to acknowledge some of the limitations of our study, but also to emphasize another point; because finding an association (or not) between rates of diversification can occur irrespective of whether traits are functionally coupled, such findings should be interpreted with care (Uyeda et al. 2021).

With this in mind, we note that similar patterns have been observed in other empirical studies. For example, lineage diversification without associated phenotypic diversification has been reported in a diverse array of taxonomic groups, including plethodontid salamanders (Adams et al. 2009), birds (Crouch & Ricklefs, 2019), and ferns (Testo & Sundue, 2018). However, lags in the timing of phenotypic diversification (*e*.*g*., Folk et al. 2018) and niche diversification (*e*.*g*., McCormack et al. 2010) are apparently less common. This discrepancy may be explainable if the initial burst in the rate of lineage diversification is not due to increased ecological opportunity (*i*.*e*., niche-neutral), but subsequent diversification dynamics are indeed primarily density-dependent. Initial niche-neutral diversification can lead to a burst in the rate of lineage diversification without associated bursts in phenotypic or niche diversification rates (Aguilée et al. 2018; Folk et al. 2018). If, subsequent to this burst, increases in ecological opportunity enable ecological specialization and adaptation to novel environments, ecologically-driven diversification processes (*e*.*g*., niche partitioning and divergence) can begin to supersede niche-neutral processes, leading to accelerations in phenotypic and niche diversification (Aguilée et al. 2018). Indeed, this logic was used to explain lags in phenotypic and niche diversification in the Saxifragales (Folk et al. 2018). Contrary, to our findings, however, Folk et al. (2018) did not observe a slowdown in Saxifragales lineage diversification rates. This was attributed to continued formation and expansion of novel habitat, consistent with ecological opportunity caused from climate cooling and aridification after the mid-Miocene Climatic Optimum. While a slowdown is clearly evident in *Penstemon* (Figure 3), this is not in violation of theoretical expectations for density-dependent diversification dynamics; interspecific competition accelerates trait diversity, but not necessarily species richness, and in fact, competition can cause a slowdown in rates of lineage diversification even when there are no explicit ecological limits (Aristide & Morlon, 2019). Overall, our findings are consistent with initial niche-neutral diversification in *Penstemon*, followed by density-dependent diversification processes that simultaneously increased phenotypic diversity and led to a slowdown in lineage diversification rates.

### Geography, ecological opportunity, and the primary mode of speciation

Although the initial burst in lineage diversification rates in *Penstemon* is consistent with niche-neutral diversification, we did also find evidence of a link between lineage diversification rate and ecological opportunity in the form of global temperature (Figure 4; Supplemental Table 6). Relatedly, we found evidence for higher rates of lineage diversification in warmer environments and in larger plants (Table 4). If the initial burst in lineage diversification is primarily niche-neutral, why do there still appear to be ecological predictors of lineage diversification rates in *Penstemon*?

The timing of the burst in *Penstemon* lineage diversification rate corresponds to the onset of the Quaternary Period approximately 2.58 MYA (Head & Gibbard, 2015). With the Quaternary Period came severe environmental changes, including the development of ice sheets and the intensification of glaciation in the Northern Hemisphere (Head & Gibbard, 2015). A natural conclusion, then, would be to associate the shift in the rate of lineage diversification in *Penstemon* with the increased ecological opportunity brought about by the cooling of the climate and the appearance of novel niches due to Quaternary glaciation. However, it is perhaps more likely that these relationships are a byproduct of the timing of the *Penstemon* radiation, and not a cause, *per se*. While ecological opportunity does increase during phases of rapid environmental change (Wellborn & Langerhans, 2015), and undoubtedly, Quaternary glaciation had profound effects on the geographic distributions and genomes of species (Hewitt, 2000), the lack of corresponding phenotypic and niche diversification in *Penstemon* suggests that although initial lineage diversification coincided with large-scale environmental change, speciation was still mostly occurring in environmental conditions similar to ancestral habitats. In this scenario, initial lineage diversification in *Penstemon* may have been a primarily geographic phenomenon: spurred by environmental changes, but only because those changes resulted in the formation of abundant habitat similar to habitats in which *Penstemon* species were already found.

Evidence from species distribution models in *Penstemon* subgenus *Dasanthera* lends some support for this scenario (Stone & Wolfe, 2021). For subgenus *Dasanthera*, which is sister to the rest of *Penstemon* and is not included in the lineage diversification rate shift, periods of peak glacial expanse likely brought about large increases in suitable habitat, particularly in the Columbia Basin and the Great Basin (Stone & Wolfe, 2021). These geographic regions, while ecologically similar to ancestral habitats, are now disjunct from current species’ distributions, although a few narrowly endemic taxa (*i*.*e*., *P. davidsonii* var. *praeteritus* and *P. fruticosus* var. *serratus*) still persist in isolated mountain ranges. A similar scenario for other *Penstemon* species, then, seems plausible, especially given the abundance of narrowly endemic species (most of which, consequently, are found in the Great Basin) and the lack of signal for shifts in niche diversification in our analyses (Supplemental Table 5; Supplemental Figure 1). Indeed, a strong geographic component to speciation has been confirmed to be an important driver of diversification in *Penstemon*, with associations between founder-event speciation events and elevated rates of net lineage diversification pointing to a likely mechanism for the rapid radiation of the genus (Wolfe et al. 2021). The importance of these founder-event speciation events suggests that the dispersal of *Penstemon* species to previously unoccupied areas was key to the diversification of the genus (Wolfe et al. 2021). Likewise, the timing of these diversification bursts, coincident with pulses of glaciation during the Pleistocene, likely reflect repeated migration events into the Intermountain Region (Cronquist, 1978), and are consistent with expectations for species’ responses to ecological opportunity in an adaptive radiation (Simpson, 1953; Wolfe et al. 2021). With respect to the mechanism of speciation, a primarily niche-neutral process with a strong role of geography could be responsible for such patterns of lineage diversification. One possible example is the process of ‘budding’ speciation (*sensu* Anacker & Strauss, 2014), whereby larger-ranged progenitor species give rise to narrow-ranged species, often causing closely related species to be sympatric. Density-dependent processes could then follow this initial niche-neutral diversification, resulting in a slowdown of the overall lineage diversification process, increasing clade-specific rates of phenotypic diversification, and generating associations between species’ phenotypes and the environment in which they live. Future studies on the geography of speciation in *Penstemon*, especially regarding asymmetry in the size of species’ distributions and the degree of ecological and phenotypic overlap between close relatives, would be able to assess the support for this hypothesis.

Despite the apparent importance of niche-neutral processes in generating diversity in *Penstemon*, the influence of adaptive processes, particularly those related to floral evolution and the selective pressures induced by pollinators, should not be ignored. While the formation of polyploid taxa is certainly an important diversification mechanism (*e*.*g*., Keck, 1932, 1945), with virtually every *Penstemon* researcher since Pennell acknowledging the incredible diversity of floral shapes, sizes, and colors, it is clear that pollinator pressure has played a substantial role in the evolutionary history of this genus (Pennell, 1935; Straw, 1966; Wilson et al. 2004; Wolfe et al. 2006; Wessinger et al. 2019). Perhaps unsurprisingly, then, many studies have provided evidence of adaptation to different pollinators in *Penstemon* (*e*.*g*., Straw, 1956, 1963; Crosswhite & Crosswhite, 1966), with transitions from insect- to hummingbird pollination especially common (Straw, 1963; Wilson et al. 2007; Wolfe et al. 2006, 2021). It is for this reason that the most likely density-dependent processes driving post-dispersal speciation to unoccupied niches involve adaptations to different pollinator types. This speciation process – relatively sluggish, compared to the multitude of founder-event speciation events associated with dispersal to unoccupied niches – would account for both the slowdown in lineage diversification rates (Figure 3) and clade-specific bursts in phenotypic diversification rates (Supplemental Figure 1) we have presented here. Supporting this claim, transitions from insect- to hummingbird pollination, while relatively common in *Penstemon*, have been associated with slowdowns in rates of diversification (Wessinger et al. 2019).

In light of this, it is important to consider the limitations of the analyses we have conducted in this study. By virtue of reducing the dimensionality of our data set into a single summary statistic, we capture less of the variance between species than is explained by the data set as a whole. By considering only the first principal component for niche and phenotype diversification as univariate traits, our analyses represent a biased sample of what is truly a multivariate pattern (Uyeda et al. 2015). While reducing the complexity of the data in this way is necessary to perform many of the analyses we have presented here, it also creates a dilemma we must reconcile; it is possible – perhaps even likely – that the summary statistics employed in this study, despite explaining a large proportion of variance, may not be biologically meaningful summarizations of phenotypic and niche diversity in *Penstemon*. Future comparative studies interested in explaining diversification patterns in *Penstemon* may therefore benefit from generating a phenotypic data set more acutely focused on explaining the diversity of floral types evident in the genus.

