## supplement for "Asynchronous rates of lineage, phenotype, and niche diversification in a continental-scale adaptive radiation"

Table of Contents:


| | |
|---|---|
| Supplemental Figure 1 | Page 2 |
| Supplemental Table 1 | Page 3 |
| Supplemental Table 2 | Page 4 |
| Supplemental Table 3 | Page 5 |
| Supplemental Table 4 | Pages 6-9 |
| Supplemental Table 5 | Pages 10-11 |

*Supplemental Figure 1*. Clade-specific rates of phenotype (left), and niche (right) diversification through time. Clades differing from the general pattern of decreasing diversification rates are indicated in the top-left corner.

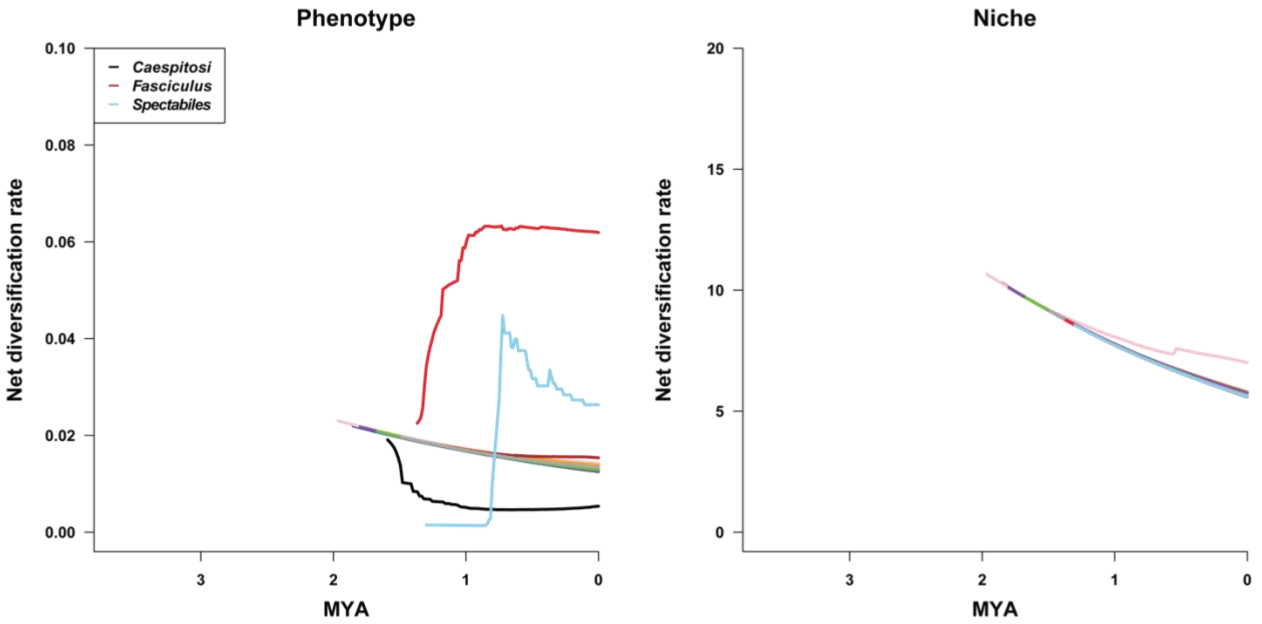

*Supplemental Table 1.* Species with data from additional sources

| Species | Occurrence points | Herbarium specimens |
|---|---|---|
| *P. amphorellae* | - | 1,2,3,4 |
| *P. campanulatus* | - | 1,5 |
| *P. coriaceus* | y | - |
| *P. fasciculatus* | - | 2,6,7 |
| *P. gentianoides* | - | 1,8 |
| *P. goodrichii* | y | - |
| *P. hartwegii* | - | 1 |
| *P. hesperius* | y | - |
| *P. hidalgensis* | - | 2 |
| *P. imberbis* | - | 2 |
| *P. isophyllus* | - | 1,2 |
| *P. leonensis* | - | 1,2,8 |
| *P. miniatus* | - | 2,4,7 |
| *P. occiduus* | - | 9 |
| *P. pahutensis* | y | - |
| *P. parvus* | y | - |
| *P. penlandii* | y | - |
| *P. plagapineus* | - | 2,10 |
| *P. reidmoranii* | y | - |
| *P. roseus* | - | 1,10 |
| *P. saltarius* | - | 2 |
| *P. smallii* | y | - |
| *P. versicolor* | y | - |
| *P. vizcainensis* | - | 2 |
| *P. wislizeni* | y | 7,11 |

1, Arizona State University Vascular Plant Herbarium
2, Steere Herbarium at the New York Botanical Garden
3, Utah State University Intermountain Herbarium
4, National Autonomous University of Mexico Herbarium
5, University of Colorado Museum Herbarium
6, Desert Botanical Garden Herbarium
7, California Botanic Garden Herbarium
8, Harvard University Herbaria and Botanical Museum
9, United States National Herbarium
10, John G. Searle Herbarium at the Field Museum of Natural History
11, Deaver Herbarium at Norther Arizona University

*Supplemental Table 2.* Phenotypic traits collected for all species

| Trait |
|---|
| Height (cm) |
| Leaf length (mm) |
| Leaf width (mm) |
| Corolla length (mm) |
| Corolla tube length (mm) |
| Corolla Throat diameter (mm) |
| Staminode length (mm) |
| Style length (mm) |
| Form |
| Basal leaves |
| Stamen location |
| Staminode location |
| Lifespan |
| Inflorescence |
| Flower presentation |
| Flower shape |
| Corolla throat expansion |
| Corolla lower topography |
| Anther dehiscence pattern |

*Supplemental Table 3.* Bayes Factors and posterior probabilities of *BAMM* lineage diversification

| Bayes Factors | | | |
|---|---|---|---|
| Number of shifts | 1 | 2 | 3 |
| 1 | - | 6.013 | 34.407 |
| 2 | 0.166 | - | 5.722 |
| 3 | 0.029 | 0.175 | - |
| Posterior Probabilities | 0.917 | 0.076 | 0.006 |

*Supplemental Table 4.* Results of *BAMM* phenotype diversification

| | Bayes Factors | | | | | | |
|---|---|---|---|---|---|---|---|
| Number of shifts | 2 | 3 | 4 | 5 | 6 | 7 | 8 |
| 2 | - | 1.9E-01 | 5.0E-02 | 5.1E-03 | 7.2E-04 | 1.7E-04 | 6.1E-05 |
| 3 | 5.3E+00 | - | 2.7E-01 | 2.7E-02 | 3.9E-03 | 9.3E-04 | 3.2E-04 |
| 4 | 2.0E+01 | 3.8E+00 | - | 1.0E-01 | 1.4E-02 | 3.5E-03 | 1.2E-03 |
| 5 | 1.9E+02 | 3.7E+01 | 9.7E+00 | - | 1.4E-01 | 3.4E-02 | 1.2E-02 |
| 6 | 1.4E+03 | 2.6E+02 | 6.9E+01 | 7.1E+00 | - | 2.4E-01 | 8.4E-02 |
| 7 | 5.7E+03 | 1.1E+03 | 2.9E+02 | 2.9E+01 | 4.1E+00 | - | 3.5E-01 |
| 8 | 1.6E+04 | 3.1E+03 | 8.2E+02 | 8.5E+01 | 1.2E+01 | 2.9E+00 | - |
| 9 | 4.4E+04 | 8.3E+03 | 2.2E+03 | 2.3E+02 | 3.2E+01 | 7.7E+00 | 2.7E+00 |
| 10 | 9.3E+04 | 1.7E+04 | 4.7E+03 | 4.8E+02 | 6.7E+01 | 1.6E+01 | 5.7E+00 |
| 11 | 1.8E+05 | 3.5E+04 | 9.2E+03 | 9.5E+02 | 1.3E+02 | 3.2E+01 | 1.1E+01 |
| 12 | 3.3E+05 | 6.2E+04 | 1.7E+04 | 1.7E+03 | 2.4E+02 | 5.8E+01 | 2.0E+01 |
| 13 | 5.6E+05 | 1.0E+05 | 2.8E+04 | 2.9E+03 | 4.0E+02 | 9.8E+01 | 3.4E+01 |
| 14 | 9.4E+05 | 1.8E+05 | 4.7E+04 | 4.8E+03 | 6.8E+02 | 1.6E+02 | 5.7E+01 |
| 15 | 1.4E+06 | 2.6E+05 | 6.9E+04 | 7.0E+03 | 9.9E+02 | 2.4E+02 | 8.3E+01 |
| 16 | 2.0E+06 | 3.8E+05 | 1.0E+05 | 1.0E+04 | 1.5E+03 | 3.5E+02 | 1.2E+02 |
| 17 | 2.8E+06 | 5.3E+05 | 1.4E+05 | 1.4E+04 | 2.0E+03 | 4.9E+02 | 1.7E+02 |
| 18 | 3.8E+06 | 7.2E+05 | 1.9E+05 | 2.0E+04 | 2.8E+03 | 6.7E+02 | 2.3E+02 |
| 19 | 5.8E+06 | 1.1E+06 | 2.9E+05 | 3.0E+04 | 4.2E+03 | 1.0E+03 | 3.5E+02 |
| 20 | 6.7E+06 | 1.3E+06 | 3.4E+05 | 3.5E+04 | 4.9E+03 | 1.2E+03 | 4.1E+02 |
| 21 | 8.9E+06 | 1.7E+06 | 4.5E+05 | 4.6E+04 | 6.5E+03 | 1.6E+03 | 5.4E+02 |
| 22 | 1.3E+07 | 2.4E+06 | 6.5E+05 | 6.6E+04 | 9.4E+03 | 2.3E+03 | 7.9E+02 |
| 23 | 1.5E+07 | 2.9E+06 | 7.7E+05 | 7.9E+04 | 1.1E+04 | 2.7E+03 | 9.3E+02 |
| 24 | 1.5E+07 | 2.9E+06 | 7.7E+05 | 7.9E+04 | 1.1E+04 | 2.7E+03 | 9.3E+02 |
| 25 | 1.4E+07 | 2.6E+06 | 7.0E+05 | 7.2E+04 | 1.0E+04 | 2.4E+03 | 8.5E+02 |
| 26 | 1.1E+07 | 2.1E+06 | 5.6E+05 | 5.7E+04 | 8.1E+03 | 2.0E+03 | 6.8E+02 |
| 27 | 3.4E+07 | 6.3E+06 | 1.7E+06 | 1.7E+05 | 2.4E+04 | 5.9E+03 | 2.0E+03 |
| 28 | 4.5E+07 | 8.4E+06 | 2.2E+06 | 2.3E+05 | 3.2E+04 | 7.8E+03 | 2.7E+03 |
| Posterior Probabilities | 0.00 | 0.00 | 0.00 | 0.01 | 0.03 | 0.06 | 0.09 |

| 9 | 10 | 11 | 12 | 13 | 14 | 15 | 16 | 17 |
|---|---|---|---|---|---|---|---|---|
| 2.3E-05 | 1.1E-05 | 5.4E-06 | 3.0E-06 | 1.8E-06 | 1.1E-06 | 7.3E-07 | 4.9E-07 | 3.5E-07 |
| 1.2E-04 | 5.7E-05 | 2.9E-05 | 1.6E-05 | 9.6E-06 | 5.7E-06 | 3.9E-06 | 2.6E-06 | 1.9E-06 |
| 4.5E-04 | 2.1E-04 | 1.1E-04 | 6.0E-05 | 3.6E-05 | 2.1E-05 | 1.5E-05 | 9.9E-06 | 7.1E-06 |
| 4.4E-03 | 2.1E-03 | 1.1E-03 | 5.9E-04 | 3.5E-04 | 2.1E-04 | 1.4E-04 | 9.6E-05 | 6.9E-05 |
| 3.1E-02 | 1.5E-02 | 7.5E-03 | 4.2E-03 | 2.5E-03 | 1.5E-03 | 1.0E-03 | 6.8E-04 | 4.9E-04 |
| 1.3E-01 | 6.1E-02 | 3.1E-02 | 1.7E-02 | 1.0E-02 | 6.1E-03 | 4.2E-03 | 2.8E-03 | 2.0E-03 |
| 3.7E-01 | 1.8E-01 | 8.9E-02 | 5.0E-02 | 3.0E-02 | 1.8E-02 | 1.2E-02 | 8.1E-03 | 5.8E-03 |
| - | 4.7E-01 | 2.4E-01 | 1.3E-01 | 7.9E-02 | 4.7E-02 | 3.2E-02 | 2.2E-02 | 1.6E-02 |
| 2.1E+00 | - | 5.1E-01 | 2.8E-01 | 1.7E-01 | 9.9E-02 | 6.8E-02 | 4.6E-02 | 3.3E-02 |
| 4.2E+00 | 2.0E+00 | - | 5.5E-01 | 3.3E-01 | 2.0E-01 | 1.3E-01 | 9.1E-02 | 6.5E-02 |
| 7.5E+00 | 3.6E+00 | 1.8E+00 | - | 6.0E-01 | 3.5E-01 | 2.4E-01 | 1.6E-01 | 1.2E-01 |
| 1.3E+01 | 6.0E+00 | 3.0E+00 | 1.7E+00 | - | 6.0E-01 | 4.1E-01 | 2.8E-01 | 2.0E-01 |
| 2.1E+01 | 1.0E+01 | 5.1E+00 | 2.8E+00 | 1.7E+00 | - | 6.8E-01 | 4.6E-01 | 3.3E-01 |
| 3.1E+01 | 1.5E+01 | 7.4E+00 | 4.1E+00 | 2.5E+00 | 1.5E+00 | - | 6.8E-01 | 4.9E-01 |
| 4.6E+01 | 2.2E+01 | 1.1E+01 | 6.1E+00 | 3.6E+00 | 2.2E+00 | 1.5E+00 | - | 7.2E-01 |
| 6.4E+01 | 3.0E+01 | 1.5E+01 | 8.5E+00 | 5.1E+00 | 3.0E+00 | 2.1E+00 | 1.4E+00 | - |
| 8.7E+01 | 4.1E+01 | 2.1E+01 | 1.2E+01 | 6.9E+00 | 4.1E+00 | 2.8E+00 | 1.9E+00 | 1.4E+00 |
| 1.3E+02 | 6.2E+01 | 3.1E+01 | 1.7E+01 | 1.0E+01 | 6.2E+00 | 4.2E+00 | 2.8E+00 | 2.0E+00 |
| 1.5E+02 | 7.2E+01 | 3.6E+01 | 2.0E+01 | 1.2E+01 | 7.2E+00 | 4.9E+00 | 3.3E+00 | 2.4E+00 |
| 2.0E+02 | 9.6E+01 | 4.8E+01 | 2.7E+01 | 1.6E+01 | 9.5E+00 | 6.5E+00 | 4.4E+00 | 3.2E+00 |
| 2.9E+02 | 1.4E+02 | 7.0E+01 | 3.9E+01 | 2.3E+01 | 1.4E+01 | 9.4E+00 | 6.4E+00 | 4.6E+00 |
| 3.5E+02 | 1.7E+02 | 8.3E+01 | 4.6E+01 | 2.8E+01 | 1.6E+01 | 1.1E+01 | 7.6E+00 | 5.5E+00 |
| 3.5E+02 | 1.7E+02 | 8.3E+01 | 4.6E+01 | 2.8E+01 | 1.6E+01 | 1.1E+01 | 7.6E+00 | 5.5E+00 |
| 3.2E+02 | 1.5E+02 | 7.6E+01 | 4.2E+01 | 2.5E+01 | 1.5E+01 | 1.0E+01 | 6.9E+00 | 5.0E+00 |
| 2.5E+02 | 1.2E+02 | 6.1E+01 | 3.4E+01 | 2.0E+01 | 1.2E+01 | 8.2E+00 | 5.5E+00 | 4.0E+00 |
| 7.6E+02 | 3.6E+02 | 1.8E+02 | 1.0E+02 | 6.0E+01 | 3.6E+01 | 2.4E+01 | 1.7E+01 | 1.2E+01 |
| 1.0E+03 | 4.8E+02 | 2.4E+02 | 1.3E+02 | 8.0E+01 | 4.8E+01 | 3.3E+01 | 2.2E+01 | 1.6E+01 |
| 0.12 | 0.12 | 0.12 | 0.11 | 0.09 | 0.08 | 0.06 | 0.04 | 0.03 |

| 18 | 19 | 20 | 21 | 22 | 23 | 24 | 25 | 26 |
|---|---|---|---|---|---|---|---|---|
| 2.6E-07 | 1.7E-07 | 1.5E-07 | 1.1E-07 | 7.7E-08 | 6.5E-08 | 6.5E-08 | 7.2E-08 | 8.9E-08 |
| 1.4E-06 | 9.2E-07 | 7.9E-07 | 6.0E-07 | 4.1E-07 | 3.5E-07 | 3.5E-07 | 3.8E-07 | 4.8E-07 |
| 5.2E-06 | 3.5E-06 | 3.0E-06 | 2.2E-06 | 1.5E-06 | 1.3E-06 | 1.3E-06 | 1.4E-06 | 1.8E-06 |
| 5.1E-05 | 3.4E-05 | 2.9E-05 | 2.2E-05 | 1.5E-05 | 1.3E-05 | 1.3E-05 | 1.4E-05 | 1.7E-05 |
| 3.6E-04 | 2.4E-04 | 2.1E-04 | 1.5E-04 | 1.1E-04 | 9.0E-05 | 9.0E-05 | 9.9E-05 | 1.2E-04 |
| 1.5E-03 | 9.9E-04 | 8.5E-04 | 6.4E-04 | 4.4E-04 | 3.7E-04 | 3.7E-04 | 4.1E-04 | 5.1E-04 |
| 4.3E-03 | 2.9E-03 | 2.4E-03 | 1.8E-03 | 1.3E-03 | 1.1E-03 | 1.1E-03 | 1.2E-03 | 1.5E-03 |
| 1.2E-02 | 7.7E-03 | 6.6E-03 | 5.0E-03 | 3.4E-03 | 2.9E-03 | 2.9E-03 | 3.2E-03 | 4.0E-03 |
| 2.4E-02 | 1.6E-02 | 1.4E-02 | 1.0E-02 | 7.2E-03 | 6.1E-03 | 6.1E-03 | 6.7E-03 | 8.3E-03 |
| 4.8E-02 | 3.2E-02 | 2.7E-02 | 2.1E-02 | 1.4E-02 | 1.2E-02 | 1.2E-02 | 1.3E-02 | 1.6E-02 |
| 8.6E-02 | 5.8E-02 | 4.9E-02 | 3.7E-02 | 2.6E-02 | 2.2E-02 | 2.2E-02 | 2.4E-02 | 3.0E-02 |
| 1.5E-01 | 9.7E-02 | 8.3E-02 | 6.3E-02 | 4.3E-02 | 3.6E-02 | 3.6E-02 | 4.0E-02 | 5.0E-02 |
| 2.4E-01 | 1.6E-01 | 1.4E-01 | 1.1E-01 | 7.2E-02 | 6.1E-02 | 6.1E-02 | 6.7E-02 | 8.4E-02 |
| 3.6E-01 | 2.4E-01 | 2.0E-01 | 1.5E-01 | 1.1E-01 | 8.9E-02 | 8.9E-02 | 9.8E-02 | 1.2E-01 |
| 5.3E-01 | 3.5E-01 | 3.0E-01 | 2.3E-01 | 1.6E-01 | 1.3E-01 | 1.3E-01 | 1.4E-01 | 1.8E-01 |
| 7.3E-01 | 4.9E-01 | 4.2E-01 | 3.2E-01 | 2.2E-01 | 1.8E-01 | 1.8E-01 | 2.0E-01 | 2.5E-01 |
| - | 6.7E-01 | 5.7E-01 | 4.3E-01 | 3.0E-01 | 2.5E-01 | 2.5E-01 | 2.8E-01 | 3.4E-01 |
| 1.5E+00 | - | 8.6E-01 | 6.5E-01 | 4.5E-01 | 3.8E-01 | 3.8E-01 | 4.1E-01 | 5.2E-01 |
| 1.8E+00 | 1.2E+00 | - | 7.5E-01 | 5.2E-01 | 4.4E-01 | 4.4E-01 | 4.8E-01 | 6.0E-01 |
| 2.3E+00 | 1.5E+00 | 1.3E+00 | - | 6.9E-01 | 5.8E-01 | 5.8E-01 | 6.4E-01 | 8.0E-01 |
| 3.4E+00 | 2.2E+00 | 1.9E+00 | 1.5E+00 | - | 8.4E-01 | 8.4E-01 | 9.2E-01 | 1.2E+00 |
| 4.0E+00 | 2.7E+00 | 2.3E+00 | 1.7E+00 | 1.2E+00 | - | 1.0E+00 | 1.1E+00 | 1.4E+00 |
| 4.0E+00 | 2.7E+00 | 2.3E+00 | 1.7E+00 | 1.2E+00 | 1.0E+00 | - | 1.1E+00 | 1.4E+00 |
| 3.6E+00 | 2.4E+00 | 2.1E+00 | 1.6E+00 | 1.1E+00 | 9.1E-01 | 9.1E-01 | - | 1.3E+00 |
| 2.9E+00 | 1.9E+00 | 1.7E+00 | 1.3E+00 | 8.6E-01 | 7.3E-01 | 7.3E-01 | 8.0E-01 | - |
| 8.7E+00 | 5.8E+00 | 5.0E+00 | 3.8E+00 | 2.6E+00 | 2.2E+00 | 2.2E+00 | 2.4E+00 | 3.0E+00 |
| 1.2E+01 | 7.8E+00 | 6.6E+00 | 5.0E+00 | 3.5E+00 | 2.9E+00 | 2.9E+00 | 3.2E+00 | 4.0E+00 |
| 0.02 | 0.01 | 0.01 | 0.01 | 0.00 | 0.00 | 0.00 | 0.00 | 0.00 |

| 27 | 28 |
|---|---|
| 3.0E-08 | 2.2E-08 |
| 1.6E-07 | 1.2E-07 |
| 6.0E-07 | 4.5E-07 |
| 5.8E-06 | 4.4E-06 |
| 4.1E-05 | 3.1E-05 |
| 1.7E-04 | 1.3E-04 |
| 4.9E-04 | 3.7E-04 |
| 1.3E-03 | 9.9E-04 |
| 2.8E-03 | 2.1E-03 |
| 5.5E-03 | 4.1E-03 |
| 9.9E-03 | 7.4E-03 |
| 1.7E-02 | 1.2E-02 |
| 2.8E-02 | 2.1E-02 |
| 4.1E-02 | 3.1E-02 |
| 6.0E-02 | 4.5E-02 |
| 8.4E-02 | 6.3E-02 |
| 1.1E-01 | 8.6E-02 |
| 1.7E-01 | 1.3E-01 |
| 2.0E-01 | 1.5E-01 |
| 2.7E-01 | 2.0E-01 |
| 3.9E-01 | 2.9E-01 |
| 4.6E-01 | 3.4E-01 |
| 4.6E-01 | 3.4E-01 |
| 4.2E-01 | 3.1E-01 |
| 3.3E-01 | 2.5E-01 |
| - | 7.5E-01 |
| 1.3E+00 | - |
| 0.00 | 0.00 |

*Supplemental Table 5*. Results of *BAMM* niche diversification

| Bayes Factors | | | | | | | | | |
|---|---|---|---|---|---|---|---|---|---|
| Number of shifts | 0 | 1 | 2 | 3 | 4 | 5 | 6 | 7 | 8 |
| 0 | - | 0.49 | 0.33 | 0.24 | 0.19 | 0.16 | 0.13 | 0.12 | 0.10 |
| 1 | 2.05 | - | 0.67 | 0.49 | 0.39 | 0.32 | 0.26 | 0.25 | 0.22 |
| 2 | 3.06 | 1.50 | - | 0.73 | 0.58 | 0.48 | 0.38 | 0.37 | 0.32 |
| 3 | 4.22 | 2.06 | 1.38 | - | 0.80 | 0.66 | 0.53 | 0.51 | 0.44 |
| 4 | 5.28 | 2.58 | 1.72 | 1.25 | - | 0.83 | 0.66 | 0.63 | 0.55 |
| 5 | 6.35 | 3.10 | 2.07 | 1.51 | 1.20 | - | 0.79 | 0.76 | 0.67 |
| 6 | 7.99 | 3.90 | 2.61 | 1.90 | 1.51 | 1.26 | - | 0.96 | 0.84 |
| 7 | 8.33 | 4.07 | 2.72 | 1.98 | 1.58 | 1.31 | 1.04 | - | 0.87 |
| 8 | 9.53 | 4.65 | 3.11 | 2.26 | 1.80 | 1.50 | 1.19 | 1.14 | - |
| 9 | 10.83 | 5.29 | 3.53 | 2.57 | 2.05 | 1.70 | 1.36 | 1.30 | 1.14 |
| 10 | 8.95 | 4.37 | 2.92 | 2.12 | 1.69 | 1.41 | 1.12 | 1.08 | 0.94 |
| 11 | 13.33 | 6.51 | 4.35 | 3.16 | 2.52 | 2.10 | 1.67 | 1.60 | 1.40 |
| 12 | 14.99 | 7.32 | 4.89 | 3.56 | 2.84 | 2.36 | 1.88 | 1.80 | 1.57 |
| 13 | 18.32 | 8.95 | 5.98 | 4.35 | 3.47 | 2.89 | 2.29 | 2.20 | 1.92 |
| 14 | 16.66 | 8.13 | 5.44 | 3.95 | 3.15 | 2.62 | 2.08 | 2.00 | 1.75 |
| 15 | 19.99 | 9.76 | 6.52 | 4.74 | 3.78 | 3.15 | 2.50 | 2.40 | 2.10 |
| 16 | 26.65 | 13.01 | 8.70 | 6.32 | 5.04 | 4.20 | 3.34 | 3.20 | 2.80 |
| 17 | 26.65 | 13.01 | 8.70 | 6.32 | 5.04 | 4.20 | 3.34 | 3.20 | 2.80 |
| 18 | 53.30 | 26.02 | 17.40 | 12.64 | 10.09 | 8.39 | 6.67 | 6.40 | 5.60 |
| 19 | 426.42 | 208.17 | 139.18 | 101.14 | 80.71 | 67.15 | 53.37 | 51.20 | 44.77 |
| Posterior Probabilities | 0.24 | 0.25 | 0.19 | 0.13 | 0.08 | 0.05 | 0.03 | 0.02 | 0.01 |

| 9 | 10 | 11 | 12 | 13 | 14 | 15 | 16 | 17 | 18 | 19 |
|---|---|---|---|---|---|---|---|---|---|---|
| 0.09 | 0.11 | 0.08 | 0.07 | 0.05 | 0.06 | 0.05 | 0.04 | 0.04 | 0.02 | 0.00 |
| 0.19 | 0.23 | 0.15 | 0.14 | 0.11 | 0.12 | 0.10 | 0.08 | 0.08 | 0.04 | 0.00 |
| 0.28 | 0.34 | 0.23 | 0.20 | 0.17 | 0.18 | 0.15 | 0.11 | 0.11 | 0.06 | 0.01 |
| 0.39 | 0.47 | 0.32 | 0.28 | 0.23 | 0.25 | 0.21 | 0.16 | 0.16 | 0.08 | 0.01 |
| 0.49 | 0.59 | 0.40 | 0.35 | 0.29 | 0.32 | 0.26 | 0.20 | 0.20 | 0.10 | 0.01 |
| 0.59 | 0.71 | 0.48 | 0.42 | 0.35 | 0.38 | 0.32 | 0.24 | 0.24 | 0.12 | 0.01 |
| 0.74 | 0.89 | 0.60 | 0.53 | 0.44 | 0.48 | 0.40 | 0.30 | 0.30 | 0.15 | 0.02 |
| 0.77 | 0.93 | 0.63 | 0.56 | 0.45 | 0.50 | 0.42 | 0.31 | 0.31 | 0.16 | 0.02 |
| 0.88 | 1.06 | 0.71 | 0.64 | 0.52 | 0.57 | 0.48 | 0.36 | 0.36 | 0.18 | 0.02 |
| - | 1.21 | 0.81 | 0.72 | 0.59 | 0.65 | 0.54 | 0.41 | 0.41 | 0.20 | 0.03 |
| 0.83 | - | 0.67 | 0.60 | 0.49 | 0.54 | 0.45 | 0.34 | 0.34 | 0.17 | 0.02 |
| 1.23 | 1.49 | - | 0.89 | 0.73 | 0.80 | 0.67 | 0.50 | 0.50 | 0.25 | 0.03 |
| 1.38 | 1.67 | 1.13 | - | 0.82 | 0.90 | 0.75 | 0.56 | 0.56 | 0.28 | 0.04 |
| 1.69 | 2.05 | 1.38 | 1.22 | - | 1.10 | 0.92 | 0.69 | 0.69 | 0.34 | 0.04 |
| 1.54 | 1.86 | 1.25 | 1.11 | 0.91 | - | 0.83 | 0.63 | 0.63 | 0.31 | 0.04 |
| 1.85 | 2.23 | 1.50 | 1.33 | 1.09 | 1.20 | - | 0.75 | 0.75 | 0.38 | 0.05 |
| 2.46 | 2.98 | 2.00 | 1.78 | 1.45 | 1.60 | 1.33 | - | 1.00 | 0.50 | 0.06 |
| 2.46 | 2.98 | 2.00 | 1.78 | 1.45 | 1.60 | 1.33 | 1.00 | - | 0.50 | 0.06 |
| 4.92 | 5.95 | 4.00 | 3.56 | 2.91 | 3.20 | 2.67 | 2.00 | 2.00 | - | 0.13 |
| 39.38 | 47.63 | 32.00 | 28.44 | 23.27 | 25.60 | 21.33 | 16.00 | 16.00 | 8.00 | - |
| 0.01 | 0.00 | 0.00 | 0.00 | 0.00 | 0.00 | 0.00 | 0.00 | 0.00 | 0.00 | 0.00 |
